# Raman microscopy of cryofixed biological specimens for high-resolution and high-sensitivity chemical imaging

**DOI:** 10.1101/2023.11.15.567077

**Authors:** Kenta Mizushima, Yasuaki Kumamoto, Shoko Tamura, Masahito Yamanaka, Kentaro Mochizuki, Menglu Li, Syusuke Egoshi, Kosuke Dodo, Yoshinori Harada, Nicholas I. Smith, Mikiko Sodeoka, Hideo Tanaka, Katsumasa Fujita

**Affiliations:** Department of Applied Physics, Osaka University, Suita, Osaka 565-0871, Japan; Advanced Photonics and Biosensing Open Innovation Laboratory, AIST-Osaka University, Osaka University, Suita, Osaka 565-0871, Japan; Institute for Open and Transdisciplinary Research Initiatives, Osaka University, Suita, Osaka 565-0871, Japan; Department of Pathology and Cell Regulation, Kyoto Prefectural University of Medicine, Kamigyo-ku, Kyoto 602-8566, Japan; Synthetic Organic Chemistry Laboratory, RIKEN Cluster for Pioneering Research, Wako, Saitama 351-0198, Japan; Catalysis and Integrated Research Group, RIKEN Center for Sustainable Resource Science, 2-1 Hirosawa, Wako, Saitama 351-0198, Japan; Biophotonics Laboratory, Immunology Frontier Research Center, Osaka University, Suita, Osaka 565-0871, Japan

## Abstract

Raman microscopy is an emerging molecular imaging technology, yet its signal-to-noise-ratio (SNR) in measurements of biological specimens is severely limited due to the small cross-section of Raman scattering. Here, we present Raman imaging techniques of cryofixed specimens to overcome SNR limitations by enabling long exposure of specimens under highly stabilized low-temperature conditions. The observation of frozen specimens in a cryostat at a constant low temperature immediately after rapid freezing enabled the improvement of SNR and significantly enhanced spatial and spectral resolution. We also confirmed that the cryofixation can preserve physicochemical states of specimens by observing alkyne-labeled coenzyme Q in cytosol and hemeproteins in acute ischemic myocardium, which cannot be done by fixation using chemical reagents. Finally, we applied the technique for multiplex Raman imaging of label-free endogenous molecules and alkyne-tagged molecules in cryofixed HeLa cells, demonstrating its capability of high-content imaging of complex biological phenomena while maintaining physiological conditions.

## Introduction

Raman microscopy is an emerging tool for molecular analysis of biological specimens under physiological conditions [1],[2]. Distributions of biomolecules in a sample are imaged by obtaining the spatial distributions of Raman scattering, allowing the visualization of cell dynamics [3], drug effects [4],[5],[6], and cell differentiations [7],[8]. Interpreting sample-specific Raman spectra and their spatial distributions, Raman microscopy enables label-free and nondestructive classification of cells and tissues [9],[10],[11], which recently is a leading interest in the fields of biomedicine and regenerative medicine [12],[13]. The use of Raman tags has further extended Raman microscopy as a technique to track exogeneous or endogenous small molecules [14],[15]. This strategy has enabled super multiplex imaging that exploits the narrow line emission of Raman probes for simultaneous visualization of a large number of targets [16],[17].

One of the challenges in Raman microscopy is to improve the signal-to-noise ratio (SNR) in hyperspectral imaging. The intrinsic small cross-section of Raman scattering has limited the detection sensitivity and the resolutions in the spatial and spectral domains. High-intensity laser irradiation may increase the number of Raman scattering photons but can easily degrade the specimen. Using low-intensity laser irradiation can prevent alterations of biological specimens but then requires a long acquisition time in Raman imaging, leading to motion artifacts as well as physiological changes of specimens during observation. To overcome these problems, the spatial resolution and field of view (FOV) are sacrificed in a trade-off with image acquisition time [18],[19],[20]. The sample fixation by chemical reagents (*e*.*g*., paraformaldehyde (PFA)) or organic solvents has been used for suppressing the motion of biological specimen and preserving biomolecules [21],[22], but denatures cells by chemical reaction or dehydration [20],[21],[23]. Additionally, the utility of the chemical fixation is limited in fixing biological activities in motion [24], and there are molecular species that cannot be fixed by the current chemical fixation techniques [25].

Here we present a technique for high-SNR Raman imaging of biological specimens while preserving their physicochemical states under a low temperature. Cryopreservation of biological specimen was achieved by rapid freezing with liquid cryogen and subsequent cooling for low-temperature Raman observation. Long-term Raman imaging of the frozen specimen was enabled by a customized microscope cryostat that can maintain the specimen under low temperature. We confirmed the improvement of SNR with a longtime exposure without causing obvious sample photodegradation. The wide FOV and high-resolution measurement were also realized while avoiding motion blur in Raman imaging. We also confirmed that the redox state of heme proteins in a heart tissue, which changes rapidly at the physiological temperature (∼310 K), was preserved by cryofixation and visualized by low-temperature Raman imaging, demonstrating the capability of preserving chemical states of proteins.

## Results

An inverted Raman microscope equipped with a custom cryostat that enables rapid freezing and low-temperature observation was developed (Fig. **1A-B**). A specimen on a fused silica coverslip was placed on the sample mount in the cryostat, separated from the cooling block by a 20 μm height stainless spacer that prevents cell damage that would otherwise occur by its direct contact to the cooling block. The cooling block was equipped with a thermocouple, a liquid nitrogen circulation channel, and a heater for temperature control of the sample. Flow of nitrogen gas in the cryostat prevents frost generation that could occur around the sample during the cooling and observation processes.

The sample was cryofixed by rapid freezing using liquid propane at 88 K (in an amount of 40-60 ml) to avoid destruction of cell structures by large ice crystal formation [26],[27],[28]. Immediately after applying the cryogen to the sample, circulation of liquid nitrogen in the cooling block was started to stabilize the temperature of the cooling block for Raman measurement. The liquid propane remaining on the sample, that could interfere with Raman measurement, was removed via evaporation or introducing liquid nitrogen on the sample (see **Materials and Methods**). The cryostat was sealed using a sapphire glass window and a rubber ring before Raman observation. We confirmed that there was no image blurring by observation of 0.2 μm beads (Fig. **S1**), that could be potentially caused by vibration from the liquid nitrogen circulation after stabilization of the cooling block temperature.

We compared live-cell Raman images and spectra acquired under low temperature (233 K) and room temperature (293 K). The Raman images reconstructed at 750 cm^-1^ (porphyrin breathing of cytochromes), 1680 cm^-1^ (amide-I of proteins), and 2850 cm^-1^ (symmetric CH_2_ of lipids) of cryofixed cells also show the contrasts similar to those of living cells (Fig. **1C**). Although the existence of ice crystal was confirmed in the Raman spectra at the observation area (Fig. **S2A**), we did not observe destruction of cell body and intracellular organelles in a Raman image of cryofixed cells (Fig. **S2B**), which could be prevented by rapid freezing as slow freezing at a rate of 1 K/min showed significant destruction of sample structures (Fig. **S2C**). These results indicate that the ice crystals that could form during rapid freezing and subsequent temperature rise to 233 K were small enough to make morphological perturbation of the samples negligible in optical observations.

Although many cellular Raman bands appeared similarly at the low and room temperatures, the Raman peaks at 1061 and 2880 cm^-1^, which can be assigned to C−C stretching and asymmetric CH_2_ vibrational modes of lipid, respectively [29], exhibited differences between cryofixed cells and living cells (Fig. **1D**). The Raman images reconstructed from these bands (Fig. **S3A**) shows spatial distributions similar to that of lipid reconstructed from 2850 cm^-1^ (Fig. **1C**). The increases in Raman signals at 1061 and 2880 cm^-1^ can be attributed to lipid phase changes occurring at low temperature (233 K), at which the acyl chain of lipid is more ordered than at 293 K [29].

We also found that the resonant Raman scattering of intracellular molecules appeared differently at low temperatures. The Raman bands at 1153 and 1517 cm^-1^, which can be assigned to the resonant Raman scattering of carotenoids [30], appeared only at the low-temperature measurements (Fig. 1d) and showed the spatial distributions resembling those of lipid droplets (Fig. **S3B**) that can store hydrophobic vitamins [31]. The bands at 917, 968, and 1337 cm^-1^ can be assigned to cytochromes [32], and their spatial distributions were similar to cytochromes reconstructed from 750 cm^-1^ (Fig. **S3C**). The band at 1298 cm^-1^ can also be assigned to cytochromes [29],[32], while it can be assigned also to twisting deformational CH mode of lipid [29]. The spatial distribution of the 1298 cm^-1^ band was similar to mixture of those of cytochromes and lipid (Fig. **S3D**). The photobleaching effects in resonant Raman scattering from carotenoids and cytochromes [32],[33],[34] could be suppressed and appeared in the low-temperature measurement.

**Fig. 1.**
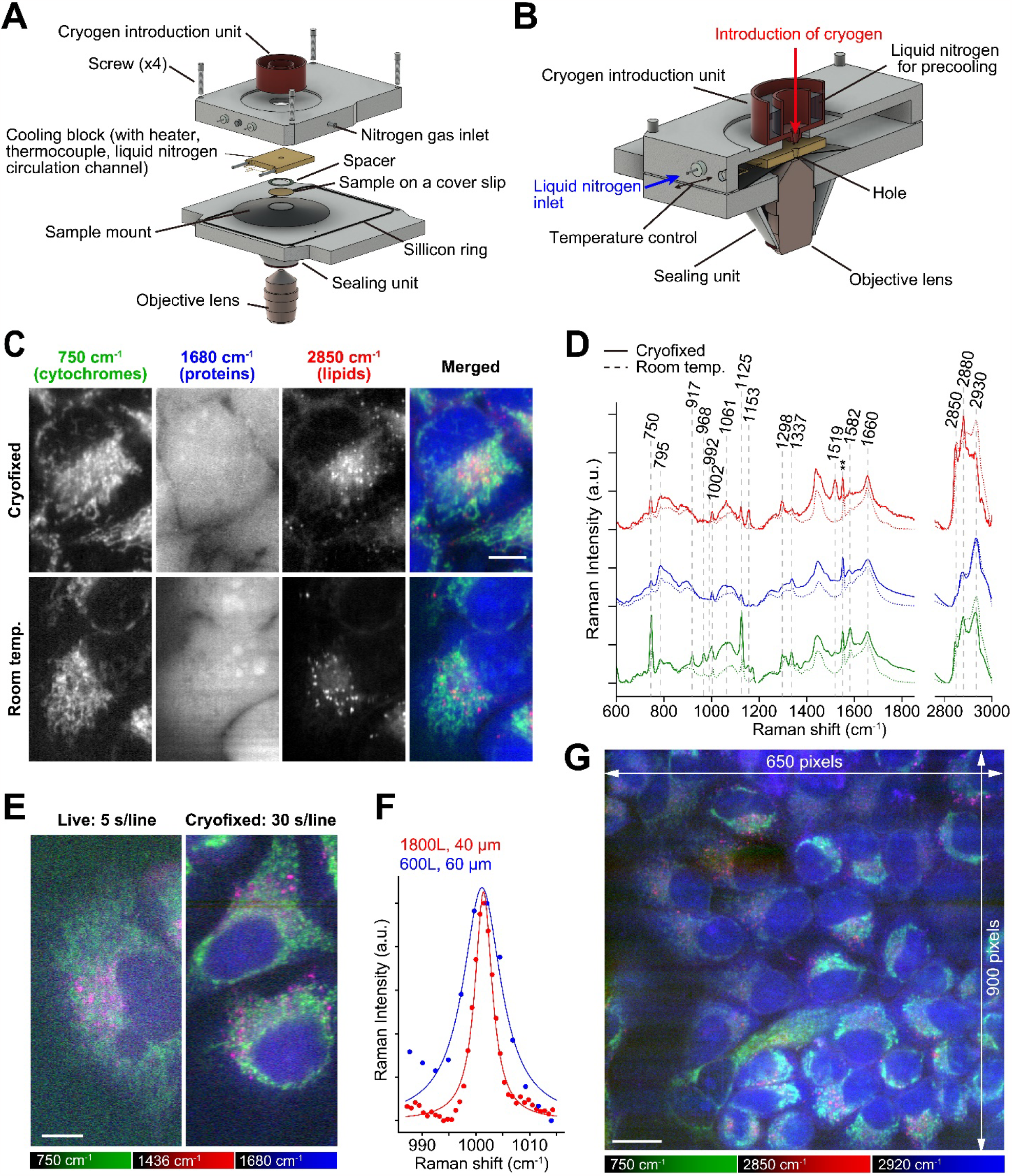
Raman imaging of cryofixed HeLa cells frozen rapidly and measured at a low temperature, enabled by development of a custom cryostat equipped with a cryogen introduction unit. (**A**) The components of the cryostat equipped with a cryogen introduction unit, together with the sample and objective lens. (**B**) The assembled cryostat. (**C**) Raman images of the rapidly frozen HeLa cells and live HeLa cells. In the spectra, atmospheric oxygen band were seen (∼1550 cm^-1^) (**). (**D**) Raman spectra of the rapidly frozen HeLa cells, acquired with the custom cryostat (at 233 K, solid lines). Room temperature (293 K) results are shown with dotted lines for comparison. The red, blue and green lines show spectra taken from the regions rich in lipids, nucleus, and cytoplasm, respectively. Spectra with different colors were vertically offset. (**E**) Raman images acquired with a high spectral resolution setting, where the slit width of the spectrophotometer was set to 40 μm (∼0.8 Airy Unit confocal slit) and the grating had a groove density of 1800 L/mm. The live and cryofixed images were acquired at 293 K and 233 K, respectively. Exposure times were 5 s/line and 30 s/line for imaging the live and cryofixed samples, respectively. (**F**) The Raman bands at 1001 cm^-1^ assigned to phenyl ring breathing of phenylalanine in HeLa cells, acquired at a high spectral resolution with a 40 μm slit width (∼0.8 Airy Unit confocal slit) and 1800 L/mm grating at cryofixed condition (red) and a low spectral resolution with a 60 μm slit width (∼1.2 Airy Unit confocal slit) and 600 L/mm grating at room temperature (blue). Red and blue solid lines are fitted using Lorentz functions. (**G**) A wide-FOV Raman image of the cryofixed HeLa cells kept at 173 K. Exposure times: 5 s/line for (**C**); 40 s/line for (**G**). Scale bars: 10 μm for (**C**) and (**E**); 20 μm for (**G**). Singular value decomposition (SVD) was applied to all data for noise reduction [35].

The high stability of sample condition is beneficial to improve the spectral and spatial resolution in Raman imaging, allowing the accumulation of Raman photons. In principle, the spectral resolution can be improved by using a spectrophotometer with higher wavelengthdispersion and a narrower entrance slit, which are not always applicable because the Raman signal decreases significantly under these conditions, especially in live-cell Raman imaging, where the SNR is typically restricted due to the limitation of exposure time to avoid the sample photodamage (Fig. **1E**). This limitation is drastically mitigated in low-temperature Raman imaging, and both high spectral and spatial resolutions can be achieved by long exposure with compensating the loss of Raman photons (Fig. **1E-F**). We confirmed the band width of 3.9 cm^-1^ at 1001 cm^-1^, assigned as phenylalanine, in the cryofixed conditions, whereas the same peak was observed to be 8.6 cm^-1^ under a practical Raman measurement condition at room temperature (Fig. **1F**).

By maintaining the low temperature of the cooling block, we can maintain the chemical conditions of the sample and the spatial distributions of biomolecules. Taking the advantage of this, we demonstrated high-resolution Raman imaging (650 x 900 image pixels) without obvious motion blur and sample drift for 10 h (Fig. **1G**). We kept the temperature of the cooling block at 173 K during measurement to avoid growth of large ice crystals in the frozen sample upon heating under prolonged laser irradiation. No obvious sample modification by laser heating was observed.

The physical fixation by rapid freezing has several advantages in comparison to chemical fixation using organic chemical reagents. As shown in Fig. **S4** the cells fixed with PFA exhibited apparent photodamage by longtime laser exposure for high SNR Raman imaging, possibly due to laser-induced heating [36] or generation of reactive oxygen species [37]. On the other hand, at the low temperature, no significant alteration in molecular distributions of the sample was confirmed after the repetitive Raman imaging (Fig. **2A**), thanks to the sample stability even under high-dose laser irradiation. We also confirmed that the Raman signal linearly increased along the exposure time (Fig. **2B**), which indicates that the biological molecules were not degraded by high-dose laser irradiation and can significantly improve the sensitivity for detecting molecules at low concentrations. We also measured 5-ethynyl-2’-deoxyuridine (EdU), an alkyne-tagged cell proliferation probe, which can be incorporated into cell nuclei, and confirmed that the distribution of EdU in cell nuclei was more clearly visualized with the longer exposure (Fig. **2C**).

Another advantage of cryofixation by rapid freezing can include instant preservation of physicochemical state of intracellular molecules under living conditions. We confirmed this advantage by observing cytochromes and alkyne-tagged coenzyme Q (AltQ2), a mobile small molecule [14], using the developed system. The PFA-fixed cells showed significant decreases in the Raman signals of cytochromes and AltQ2 (Fig. **2D**) due to oxidation of cytochromes [20] and spatial alteration of AltQ2 [38], whereas the cryofixed cells showed the distributions of cytochromes and AltQ2 similar to those in living cells (Fig. **2D**). The result demonstrates that rapid freezing and subsequently low-temperature Raman imaging can preserve and observe various cellular components at an environment close to physiological conditions, which cannot be done using conventional chemical fixation techniques.

Taking the advantages of the presented technique, we performed a high-SNR and multi-color Raman imaging of cultured cells (Fig. **2E**). In addition to Raman images of cytochromes, DNA/RNA [39], carotenoids, lipids, and proteins that are observed without labeling, three Raman tags, EdU, MitoBADY [40], and AltQ2 were simultaneously observed. We also obtained a Raman image of intracellular and extracellular water distributions, using the Raman signal that indicates ice crystal in and out of cryopreserved cells, which can be useful for understandings in the conditions of cryopreserved cells [41]. These results demonstrate the potential of cryo-Raman microscopy to detect the chemical response of endogenous molecules from their Raman spectra while using various Raman tags and probes [14],[17],[42] to perform highly multiplex observations.

A benefit of rapid freezing also lies in its ability to preserve transient biochemical reactions in biological specimens. One example is the redox state changes of cytochromes in myocardial ischemia [43]. Previous Raman imaging techniques were able to reveal the redox state change but could not visualize the spatial distribution of the redox state in the heart at a specific time point. In this research, we conducted rapid freezing and low-temperature Raman imaging of rat myocardial tissues under ischemic and non-ischemic circumstances.

We used low-temperature Raman imaging to enable redox imaging of an acute ischemic heart. A custom methodology and system were adapted for the experiment using heart tissues as follows. In this procedure, we first froze the right ventricle of myocardial tissue by liquid propane outside a cryostat (Fig. **3A**). After substituting the liquid propane with liquid nitrogen, the frozen tissue was moved into a microscope cryostat for an upright microscope (Fig. **3B**). To prevent frost formation on the sample by humid air, we immediately sealed the cryostat with the microscope objective lens and rubber sheet.

Fig. **3C** shows the results of Raman hyperspectral imaging of rat hearts rapidly frozen at 1 and 21 min after cessation of Langendorff perfusion with oxygenated Tyrode’s solution. Raman images reconstructed at 600 cm^-1^ show the distribution of reduced cytochrome *c* in cardiomyocytes. The signal intensity within the cardiomyocytes is significantly greater in the heart after 21 min of ischemia compared to 1 min, which is consistent with the findings from single-point time-lapse Raman spectroscopy [43]. The Raman spectra also reflect the trends seen in the Raman images of reduced cytochrome *c* (Fig. **3D**). Raman signals not only at 600, but also at 750 and 1128 cm^-1^ were higher in the ischemic heart than in the nonischemic heart. In addition, the Raman band at 1640 cm^-1^, identified as oxygenated myoglobin [44], showed an inverse trend to the reduced cytochrome *c* signal. Similar trends were confirmed by Raman spectroscopic measurements in live ischemic and non-ischemic rat heat tissue [45]. Our results suggest that the redox state of cytochrome *c* in cardiomyocytes was preserved by rapid freezing. In addition, we were able to acquire not only Raman spectra but also Raman images while preserving biochemical reactions by using cryofixation. The Raman image acquisition process took 27 min, during which an acutely ischemic heart would undergo changes in redox state and shape without cryofixation.

## Discussion

The combination of cryofixation and subsequent low-temperature measurement of biological specimens has overcome significant limitations in Raman microscopy, realizing a high SNR and a large FOV as well as high spatial and spectral resolution. The technique significantly reduces photodamage and motion artifacts, allowing the visualization of intracellular molecules and subcellular structures invisible in the measurement without freezing, while also enhancing quantification capabilities of Raman imaging [46],[47]. The cryofixation of chemical states, such as redox states, can further extend the advantage of Raman imaging in its capability of sensing molecular states and their environments. The extended exposure time can also achieve an even better SNR than that in coherent Raman microscopy [1].

Improving the detection sensitivity in Raman microscopy will expand the capability of optical imaging of drug substances [48],[49],[50], which is difficult with conventional fluorescence labeling. The process can also be done fully label-free, and *in vivo* applications can be realized by use of an *in vivo* rapid freezing technique [51], preserving physiological conditions of organs in living bodies and allows interrogation by Raman microscopy. In fact, Raman spectra mapping of the mouse liver frozen rapidly *in vivo* was demonstrated and showed preservation of oxygenated and deoxygenated states of hemoglobin in hepatic blood vessels [52]. In combination with this approach, our presented technique will enable high-SNR and wide-FOV Raman imaging of organs at physiological conditions at certain timepoint in living bodies with high spatial and spectral resolutions.

In our system, the detection of signals from low concentration molecules still competes with the background light that can originate from the substrates and optical components. For further improvement of detection sensitivity, reduction of such background light can be achieved by separating excitation and detection optical paths [53] as well as by use of non-emissive sample substrates [54]. Autofluorescence of the sample is also a concern in low-temperature Raman imaging as it tends to increase at lower sample temperatures and adds noise to measured Raman spectra (Fig. **S5**). Use of near-infrared or deep-ultraviolet wavelength excitation can help further improvement of SNR in low-temperature Raman imaging by eliminating the fluorescent background[55],[56].

Another potential issue in the presented method is the formation of large ice crystals and potential sample modification. In our experiments, we did not observe formation of large ice crystals during Raman observation (Fig. **S2B**). However, the Raman spectrum measurement indicated the formation of tiny ice crystals that cannot be observed by the spatial resolution of optical microscopy but may be found by higher resolution imaging techniques, such as electron microscopy and super-resolution optical microscopy. Despite the initial rapid freezing process not leading to the development of large ice crystals within cells, specifically at a cooling rate of 10^4^-10^5^ K/s [57], elevating the temperature to evaporate the liquid propane may cause the formation of large ice crystals, depending on the cell and its intracellular state [58]. In such a case, the liquid propane can be eliminated without increasing the temperature by rinsing with liquid nitrogen, as demonstrated in our experiments of wide-FOV imaging (Fig. **1G**), multi-Raman tag imaging (Fig. **2E**), and tissue measurement (Fig. **3C-D**). The impact of laser exposure might also potentially raise the sample temperature and trigger a phase transition in the ice. Indeed, we observed sample thawing with a laser fluence exceeding those demonstrated above and therefore adjusted the temperature between 153-233 K, depending on the exposure time, while maintaining the autofluorescence low enough for Raman measurement. Controlling the temperature and chemical conditions surrounding the sample will be key to expand the advantage of cryogenic Raman measurement further.

The presented method will further attract life scientists and clinicians when combined with other modalities. The presented method is highly compatible with cryo-electron microscopy [59], cryogenic super-resolution fluorescence microscopy [60], and correlative light and electron microscopy [61]. Such a combination allows correlative structural and chemical information for the precise analysis of biological samples, providing deep insights into biological phenomena from a chemical aspect that cannot be explored by conventional techniques.

## Materials and Methods

### Cell preparation

HeLa cells were cultured on fused silica coverslips with a thickness of 0.17 ± 0.02 mm and a diameter of 25 mm (Matsunami, S339882), and maintained in Dulbecco’s Modified Eagle Medium (DMEM, 043-30085, Wako-Fujifilm) supplemented with 10% fetal bovine serum (FBS, S1780-500U, Biowest) and 1% penicillin-streptomycin-glutamine solution (15140122, Gibco) for 48 h in the CO2 incubator at 310 K (37 °C). Prior to Raman measurements, the medium was replaced by Hank’s Balanced Salt Solution (HBSS, 082-08961, Wako-Fujifilm).

For chemical fixation, the cells were fixed with 4% PFA (163-20145, Wako-Fujifilm) at 293 K for 20 min. The fixed cells were rinsed three times and immersed in HBSS for Raman imaging.

For EdU measurements (Fig. **2C-E**), the cells were cultured without FBS for 24 h in a DMEM solution to synchronize cell cycle for all the cells. The medium was then replaced by a DMEM solution containing EdU at a concentration of 100 μM and the cells were incubated for 12 h (Fig. 2C) or 24 h (Fig. **2E**).

**Fig. 2.**
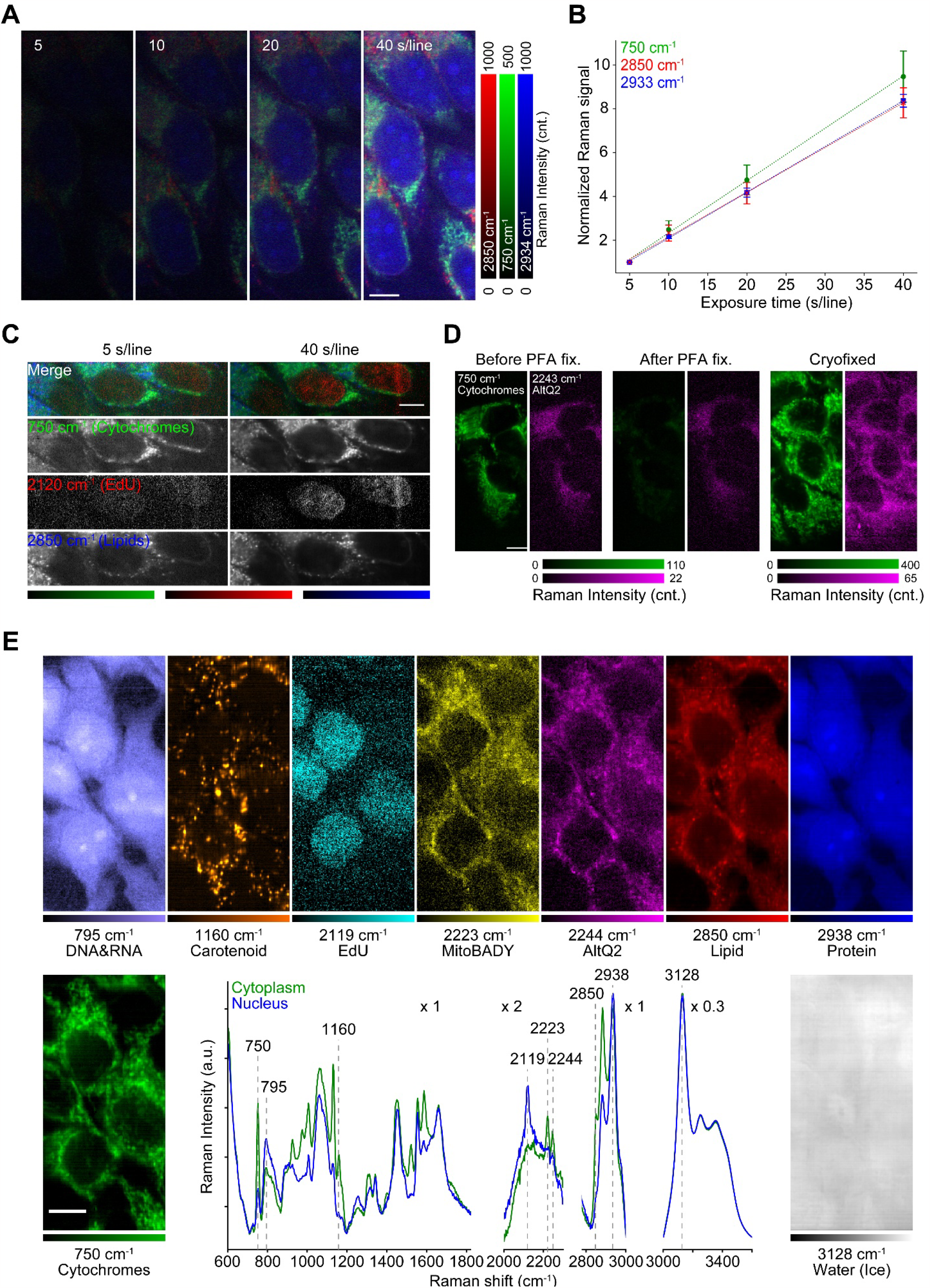
High SNR Raman imaging of HeLa cells with cryopreservation of physicochemical states. (**A**) Raman images of cryofixed HeLa cells acquired with exposure times of 5, 10, 20, and 40 s/line at 233 K. In data processing, no noise reduction was applied. (**B**) Raman signal at 750 cm^-1^, 2850 cm^-1^, and 2933 cm^-1^ of cryofixed HeLa cells to exposure time. The signal at each exposure time was normalized by the signal at 5 s/line for each Raman band. The normalized signals were averaged over 12 different regions in a. Error bars show the standard deviation for each Raman band (n = 12). The dotted lines indicate fitting curve calculated by linear functions. In data processing, no noise reduction was applied. (**C**) Raman images representing EdU incorporated in cryofixed HeLa cells, acquired with exposure times of 5 and 40 s/line at 233 K. Cytochromes and lipids are also shown to clarify the localization of EdU in nuclei. (**D**) Raman images of living cells (left), chemically fixed cells (middle), and cryofixed cells (right). Measurement temperature were 293 K (left, middle) and 233 K (right). The images represent cytochromes (green) and AltQ2 incorporated in the cells (magenta). (**E**) Nine-color Raman images and spectra measured at 203 K for the cryofixed HeLa cells loaded with EdU, MitoBADY, and AltQ2. The green, and blue lines were acquired from cytoplasm and nucleus, respectively. In the Raman spectra, the scales of lateral axes were modified in each wavenumber region, respectively. Exposure time: 10 s/line for living cells and PFA fixed cells in (**D**), 30 s/line for cryofixed cells in (**D**), 70 s/line for (**E**). Scale bars: 10 μm. SVD was applied to data shown in (**C-E**) for noise reduction [35].

**Fig. 3.**
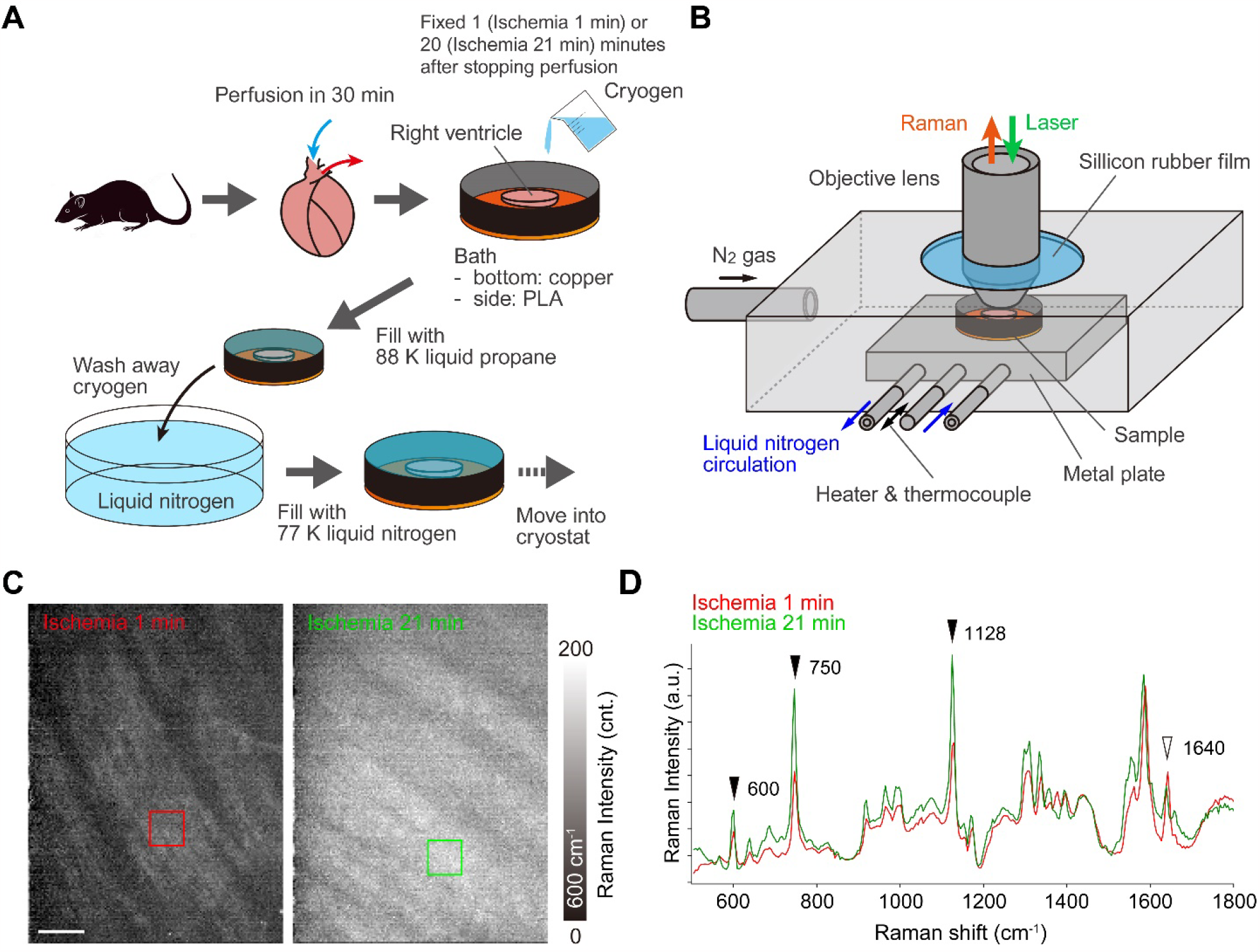
Rapid freezing and low-temperature Raman imaging of a rat heart. (**A**) Procedures of the ischemic injury, rapid freezing, and subsequent transfer of the frozen myocardial tissue to the microscope cryostat. (**B**) Schematic of the cryostat for tissue imaging by an upright Raman microscope. (**C**) Raman images reconstructed by the intensity at 600 cm^-1^, assigned to reduced cytochrome *c*. The hearts were cryofixed at 1 min (left) and 21 min (right) after cessation of perfusion and measured at 153 K. (**D**) Raman spectra of the frozen rat hearts measured at 153 K. Red and green lines exhibit average spectra taken from red and green squares in (**D**), respectively. Scale bar: 50 μm. SVD was applied to all data to separate signals from noises [35].

For AltQ2 measurements, the cells were rinsed with HBSS and incubated with AltQ2 at a concentration of 4 μM for 30 min.

For multiple Raman tag imaging, the cells were first loaded with EdU by the above-described protocol. The cells were then incubated in a DMEM solution containing MitoBADY at the concentration of 800 nM for 1 h and subsequently loaded with AltQ2 for 30 min as mentioned above.

### Tissue preparation

All of the animal experiments described in this study were conducted in accordance with the Guide for the Care and Use of Laboratory Animals (8th edition, National Academies Press, Washington DC, 2011) and with approval by the Animal Research Committee at Kyoto Prefectural University of Medicine (approval No: M2022-238). Male Wistar rats at the age of 9 weeks were purchased from Shimizu Laboratory Supplies. The animals were placed under deep general anesthesia using intraperitoneal injection with 0.1 mg/kg of medetomidine (Nippon Zenyaku Kogyo), 3.0 mg/kg of midazolam (Sandoz), and 5.0 mg/kg of butorphanol (Meiji Animal Health). After injection of heparin sodium (Mochida Pharmaceutical) into the inferior vena cava (1 U/g body weight), the hearts were quickly resected from the animals. The excised hearts were retrogradely perfused via the aorta with oxygenated Tyrode’s solution (137 mM NaCl, 4 mM KCl, 1 mM MgCl2, 0.33 mM NaH2PO4, 1.2 mM CaCl2, 10 mM HEPES, 10 mM glucose, pH = 7.4) at 310 K with a constant pressure of 9.81 kPa for approximately 3 min. Then, the retrograde perfusion with oxygenated Tyrode’s solution was switched to a constant flux rate of 10 ml/min, controlled by a micro tube pump (MP-3N, Tokyo Rikakikai). For preparation of 1-min ischemia heart, the right ventricle was cut out, placed on a copper plate, and frozen by liquid propane in 1 min after the perfusion was stopped. For preparation of 21-min ischemic heart, the whole heart was first perfused with oxygenated Tyrode’s solution for more than 30 min to stabilize its redox state and subsequently perfused with Tyrode’s solution with 20 mM 2,3-butanedione monoxime (10923-52, Nacalai Tesque) for 15 min to attenuate the contraction, according to the literature [43]. Global ischemia was then induced in the heart by stopping perfusion for 20 min [43]. Following the global ischemia, the right ventricle was cut out, placed on a copper plate, and frozen by liquid propane in 1 min.

### Slit-scanning Raman microscope equipped with a cryostat

All measurements were performed by slit-scanning Raman microscopes equipped with cryostats. For cell imaging, we set a custom cryostat equipped with a cryogen introduction unit on the sample stages of a homebuilt inverted Raman microscope system. The sample coverslip was set on the sample holder. The details of the optical setup can be seen elsewhere [2]. Briefly, a single-frequency laser at 532 nm continuous-wave oscillation (Millennia eV, Spectra Physics or Verdi, Coherent) was used for Raman excitation. The laser intensity at the sample was set at 3.0 mW/μm^2^. The laser beam was focused into a line shape by use of a cylindrical lens and focused at the sample located on an inverted microscope (Ti-E, Nikon) equipped with a 60×/0.95 NA dry objective lens (CFI Plan Apo Lambda 60XC, Nikon). The Raman scattering light generated at the sample under line illumination was collected by the same objective lens in a back scattering geometry, filtered with a longpass edge filter, and refocused at the entrance slit (60 μm) of spectrophotometer (CLP-300 or MK-300, Bunkoukeiki) equipped with a grating of 600 or 1800 L/mm. The one-dimensional distribution of Raman spectra at the sample irradiated by line illumination was then recorded by a cooled CCD camera (PIXIS 2048B or PIXIS 400BReXcelon, Teledyne Princeton Instruments). A galvanometer mirror was used to scan the sample to acquire a two-dimensional Raman hyperspectral image. The scanning pitch was set to 250 or 220 nm. In order to avoid any change in the focus position during Raman imaging, we used a focus compensation system equipped in the microscope (Nikon PFS).

For tissue imaging, we set a cryostat (THMS600, Linkam partially customized to enable the insertion of an objective lens in to the cryostat) on the sample stage of an upright slit-scanning Raman microscope commercially available (Raman-11, Nanophoton). The excitation wavelength was 532 nm and the laser intensity at the sample was set at 1.0 mW/μm^2^. The line-shaped laser beam was focused on the sample using a 20×/0.75NA dry objective lens (UPLSAP20X, Olympus) and the Raman scattering from the sample was collected using the same objective lens with a back scattering geometry. The spectrophotometer entrance slit was set at 30 μm. The grating of 300 L/mm was used. The cooled CCD camera (PIXIS 400BReXcelon, Teledyne Princeton Instruments) was used to record Raman spectra. The scanning pitch was set to 1.84 μm. The wavenumber axis of Raman spectra measurement was corrected by use of ethanol Raman bands at 434, 884,1454, and 2930 cm^-1^ as reference for cell measurements and 434, 884, 1095, and 1454 cm^-1^ as reference for tissue measurements.

### Cryofixation by liquid cryogen and subsequent cooling

Liquid propane was produced by feeding gaseous propane into a beaker cooled at 77 K by liquid nitrogen. The feeding for 4 minutes at 0.09 MPa approximately produced 60 ml of liquid propane. The liquid propane in the beaker was at around its freezing point (∼85 K) in a liquid nitrogen bath (at ∼77 K) until use for cryofixation of the sample. The cryogen temperature just before use was measured with a K-type thermocouple (RS PRO) connected to a temperature controller with a function of automatic cold junction temperature compensation (KT4R, Panasonic Industrial Devices SUNX).

For the inverted microscope system, nitrogen gas was continuously introduced into the cryostat. The sample at room temperature (293 K) was frozen by introducing the liquid propane inside the cryostat. To avoid a temperature rise of liquid propane by contact with the cooling block before reaching at the sample, the cryogen introduction unit cooled with liquid nitrogen was used to guide liquid propane to the sample through the hole (with a diameter of 2 mm) of the cooling block. The cooling block was cooled by circulating liquid nitrogen inside immediately after introduction of the cryogen until reaching at a target temperature. The introduction of nitrogen gas into the cryostat was stopped by closing the vent channel of the cryostat when the temperature of cooling block became close to the target temperature. The temperature was then stabilized by a temperature feedback control on the speed of liquid nitrogen flux. The liquid propane remaining on the sample, that could interfere with Raman measurement, was removed via evaporation by either keeping the sample temperature above the propane boiling point (231 K, for Fig. **1C-F**, and **2A-D**) or introducing liquid nitrogen (for Fig.**1G** and **2E**) on the sample to wash out the liquid propane. The temperature during Raman observation was chosen so that the autofluorescence, which can be increased at low temperature (Fig.**S5**), does not strongly disturb the Raman measurement.

For the upright microscope system, the tissue sample at room temperature (293 K) located on a copper plate with a thickness of 50 μm was frozen outside the cryostat by pouring liquid propane. The copper plate possessing the frozen sample was immediately transferred to a liquid nitrogen bath to replace the surrounding liquid propane with liquid nitrogen. The copper plate possessing the frozen sample was then quickly transferred into the cryostat that was cooled down to 153 K in advance. The liquid nitrogen surrounding the sample was evaporated naturally before starting Raman measurement. The cryostat temperature was kept through the Raman imaging by liquid nitrogen circulation in the cooling plate.

### Data processing

Acquired Raman hyperspectral image data was analyzed using a custom software developed on MATLAB environment (R2017b, Mathworks). For all the datasets, cosmic rays were removed in each CCD frame by applying a local median filter, the bias value of CCD was subtracted, and the wavenumber region of interest was cropped. Except for the quantitative SNR improvement data shown in Fig. **2A** and **2B**, each dataset was further processed as follows: SVD was applied to a Raman hyperspectral image dataset and reconstruct a new dataset for reducing the noise [35]. Raman hyperspectral data of 5 s/line and 40 s/line in Fig. **2C**, before and after PFA fixation in Fig. **2D**, were combined in each one dataset prior to SVD processing. Fluorescence background was subtracted from the Raman spectra with recursive polynomial fitting by the least-squares method [62]. To reconstruct Raman images, for all the datasets, a difference Raman intensity at two wavenumbers was mapped. All the parameters of those data processing procedures are shown in Table S1.

For the wide-field Raman image (Fig. **1E**) and multi-Raman tag imaging (Fig. **2E**), unexpected horizontal stripes appeared due to defected pixels of the spectrophotometer camera. We corrected the stripes by so called striped correction process [63], on MATLAB environment (R2023a, Mathworks) (Fig. **S6**). Briefly, each column of a reconstructed image was divided by the mean intensity projection of the image to the y axis (i.e. parallel to the line illumination) so that the stripe pattern was removed. In this process, the intensity distribution intrinsic to the sample was modified. To recover the intrinsic intensity distribution, the mean intensity projection was smoothed by a moving average filter with 51 pixels length and the derived smoothed signal was multiplied with the striped corrected image.

To quantify the signal amount improvement (Fig. **2B**), we defined a Raman signal at a wavenumber ν_1_ (Signal_ν1_) as a difference between the CCD counts at ν_1_ (I_ν1_) and another wavenumber ν_2_ (I_ν2_), where the CCD count looked similar to the baseline at ν_2_. We chose (745, 732), (2848, 2781), and (2934, 2842) cm^-1^ as (ν_1_, ν_2_) for cytochromes, lipid, and protein, respectively. We selected 12 different areas which had relatively strong Raman signals at the wavenumber of interest in Fig. **2A** (5 x 5 pixels/area for cytochromes at and lipid, 10 x 10 pixels/area for protein) and took mean values as Signal_ν1_ for each area at each exposure time by using ImageJ (1.54f). In each area, the derived Signalν1 at each exposure time was divided by that at 5 s/line for normalization. The average and standard deviation of normalized signals for 12 different areas were calculated at each exposure time. The plots of each band were fitted to a linear function by using a homemade program on Python, as the Raman signal was expected to increase linearly along exposure time on the basis of the following equation (eq. 1),

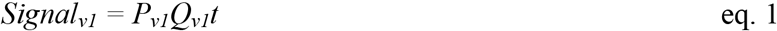

where Pv1 is the number of Raman signal photons incident to a sensor for the wavenumber ν1 per unit time, Q_ν1_ is the quantum efficiency of detector at the wavelength corresponding to the wavenumber ν1, and t is the signal accumulation time.

## Acknowledgments

The authors thank to Dr. Shogo Kawano for development of the Raman spectral imaging software.

Mr. Qiye Li and Mr. Takahiro Nagano, former members of Department of Applied Physics at Osaka

University, and Dr. Michiyo Hayakawa for helping conduction of experiments.

## Author contributions

Conceptualization: Y.K., K.F.

Methodology: K.Mi., Y.K., S.T., M.Y., K.Mo., M.L., Y.H., H.T., and K.F.

Investigation: K.Mi., Y.K., S.T., K.Mo Supervision: K.F.

Writing—original draft: K.Mi, Y.K.

Writing—review & editing: K.Mi., Y.K., M.L., N.I.S. and K.F.

## Data and materials availability

All the data used in this paper are available upon reasonable request.

