## Supplementary materials for "Raman microscopy of cryofixed biological specimens for high-resolution and high-sensitivity chemical imaging"

**This PDF file includes:**

Figs. S1 to S6

Tables S1

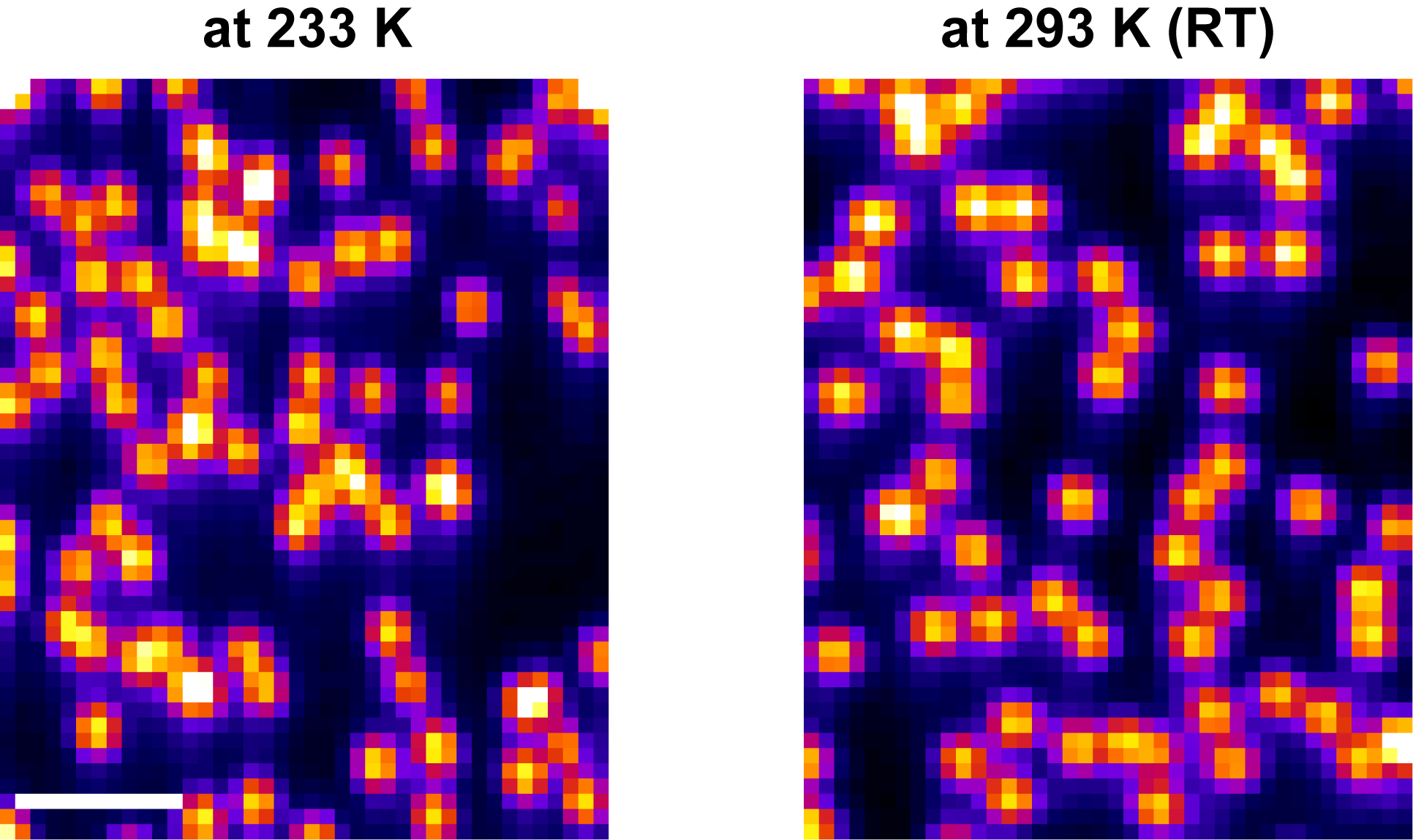

Fig. S1.

**Fluorescence images of fluorescence beads with a diameter of 0.2 µm (F8809, Invitrogen).** (Left) The beads measured at 233 K in the cryostat with liquid nitrogen circulation. (Right) The beads measured at 293 K without liquid nitrogen circulation. Scale bar: 2 µm. The fluorescence images were acquired using the slit-scanning Raman microscope described in Materials and Methods. The exposure time was 10 ms/line. The interval between acquiring neighboring lines, equivalent to the detector readout time, was 200 ms. The scanning pitch was 90 nm. The fluorescence images were reconstructed from zeroth-order light diffracted by the spectrophotometer. The slit width of spectrophotometer was 50 µm (~1.0 A.U. confocal slit).

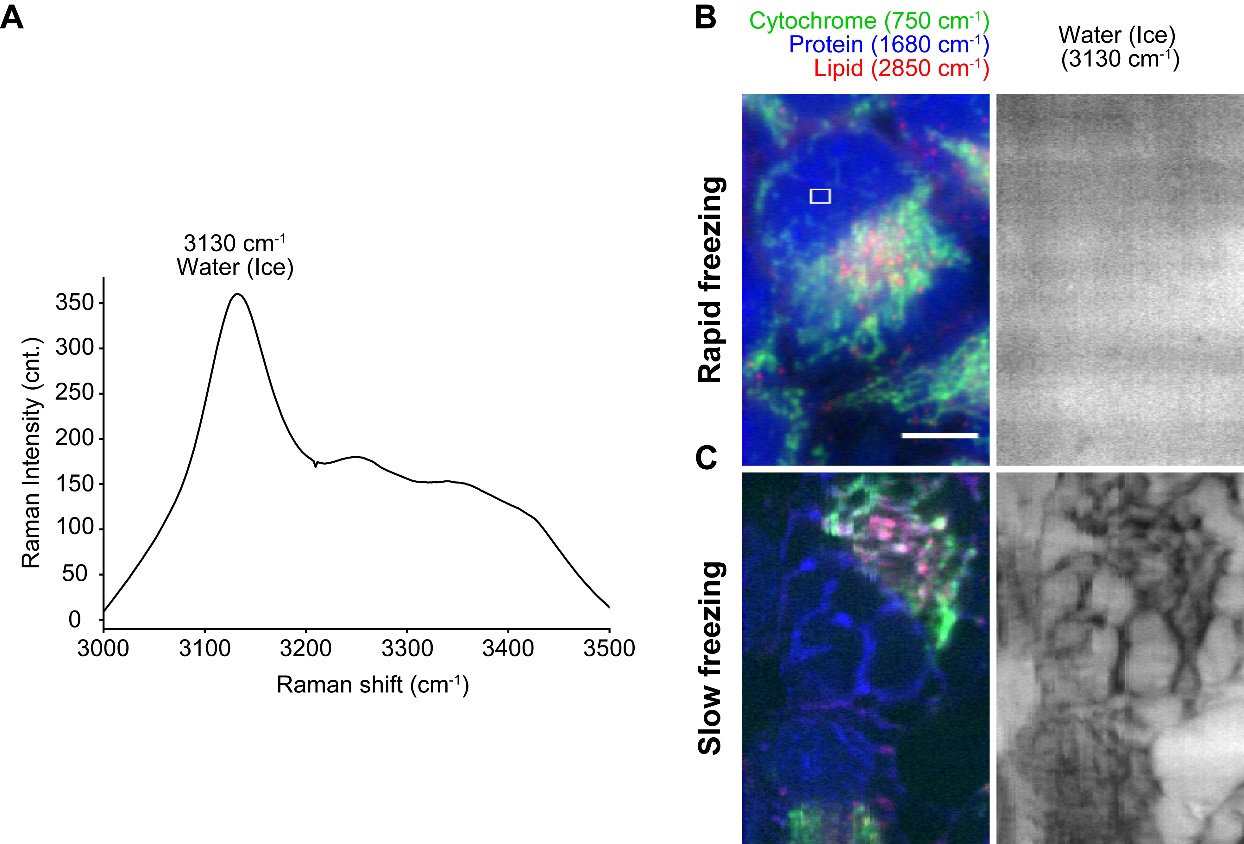

Fig. S2.

**Raman measurement of ice crystal formation in HeLa cell measurements.** (**A**) The Raman spectrum averaged over the white square in (**B**). (**B**) Raman images presenting the distributions of cytochrome, protein, and lipid in HeLa cells (left) and ice crystals over the imaging area (right), with rapid freezing by liquid propane at 88 K. (**C**) Raman images presenting the distributions of cytochrome, protein, and lipid in HeLa cells (left) and ice crystals over the imaging area (right) with slow freezing at the cooling rate of 1 K/min using the metal plate in the custom cryostat. Exposure time was 5 s/line. Scale bar: 10 µm

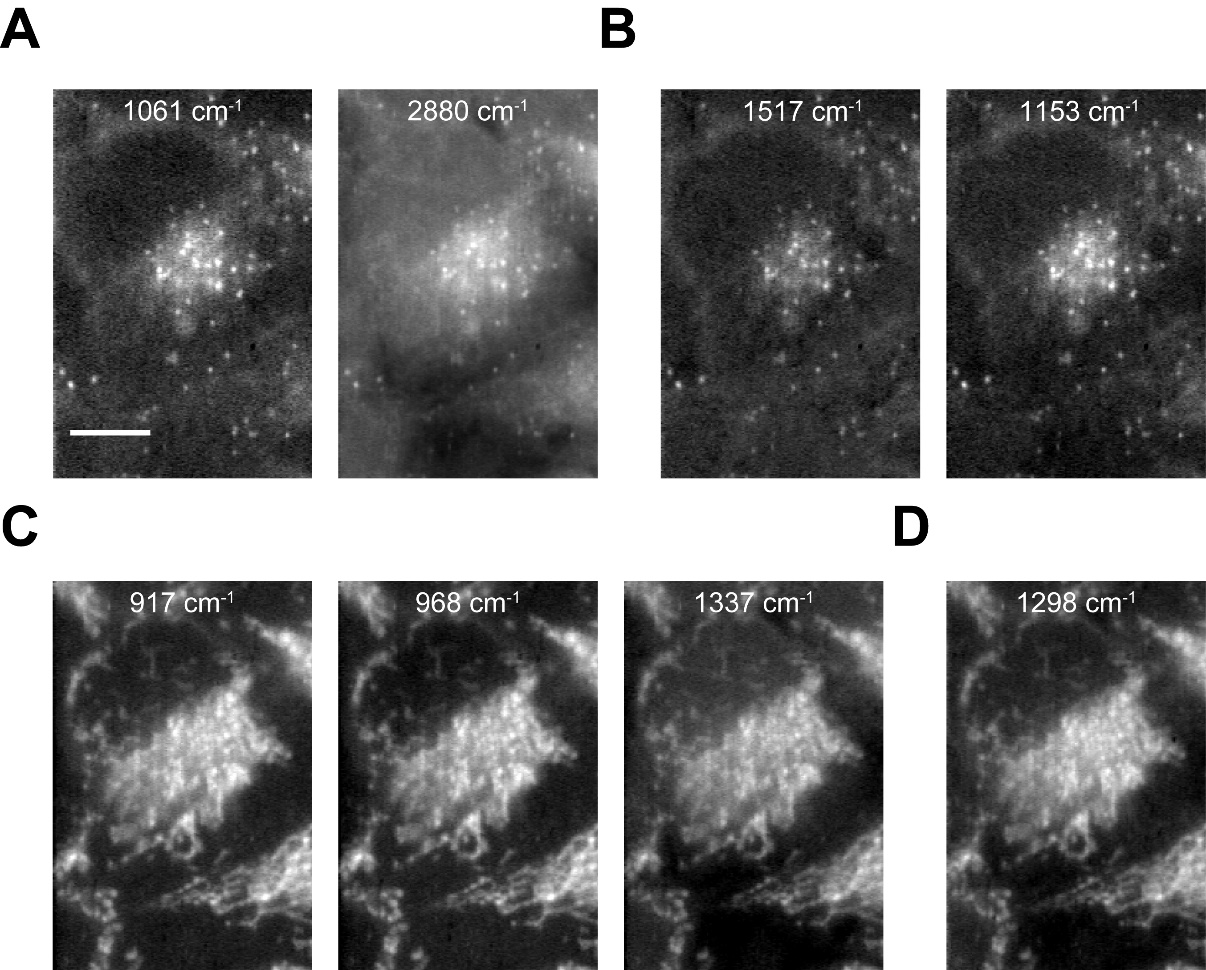

Fig. S3.

**Raman images of cryofixed HeLa cells reconstructed from Raman bands that appeared or increased only in cryofixed condition.** (**A**) The images reconstructed at 1061 and 2880 cm^-1^ (lipid) (**B**) The images reconstructed at 1151 and 1517 cm^-1^ (carotenoids). (**C**) The images reconstructed from 917, 968, and 1337 cm^-1^ (cytochromes). (**D**) The image reconstructed at 1298 cm^-1^ (cytochromes and lipid). Exposure time: 5 s/line. Scale bar: 10 µm.

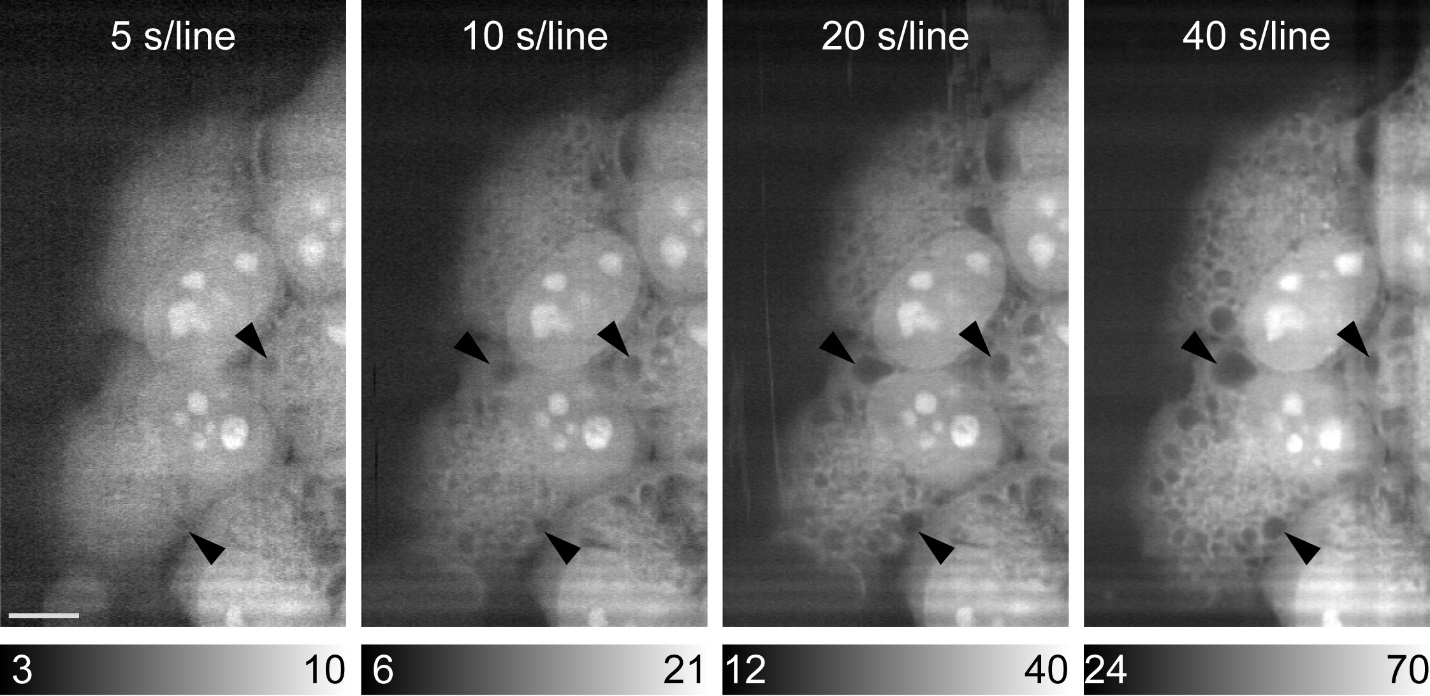

Fig. S4.

**Sample modification during repeated Raman imaging acquisitions of HeLa cells fixed by paraformaldehyde.** The images were reconstructed by the intensity at 1680 cm^-1^ assigned to the amide-I vibrational mode. Black arrows indicated the modified position during Raman imaging. The exposure times were set to 5, 10, 20, and 40 s/line for the first, second, third, and fourth acquisitions, respectively. Scale bar: 10 µm.

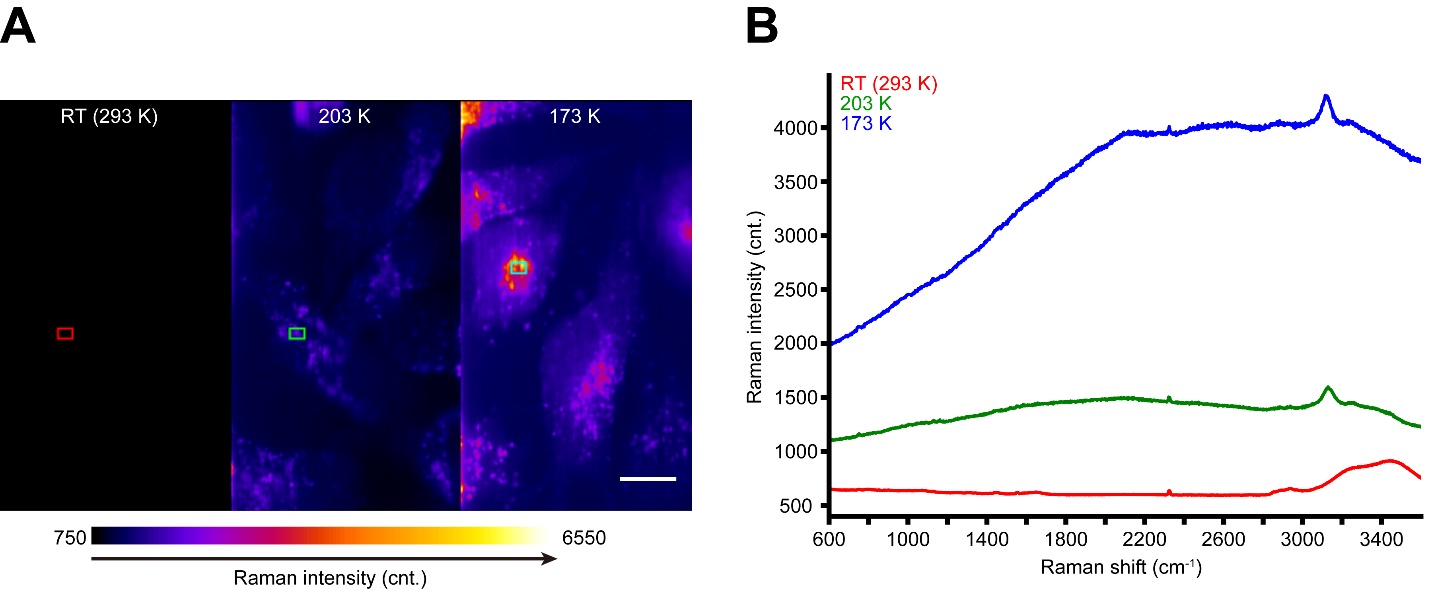

Fig. S5.

**Temperature dependence of autofluorescence in Raman imaging of cells.** (**A**) Raman images of HeLa cells at room temperature (293 K) (left), 203 K (middle), and 173 K (right) reconstructed by the intensity at 1888-2796 cm^-1^. (**B**) Average Raman spectra of HeLa cells. Red, green, and blue lines were taken at room temperature (293 K), 203 K, and 173 K, respectively. The area used for averaging spectra were framed by red, green, and cyan squares (10 pixels x 10 pixels) in a. In the data processing, no noise reduction and background subtraction was applied. Exposure time: 5 s/line. The scanning pitch was 282 nm. Scale bar: 10 µm.

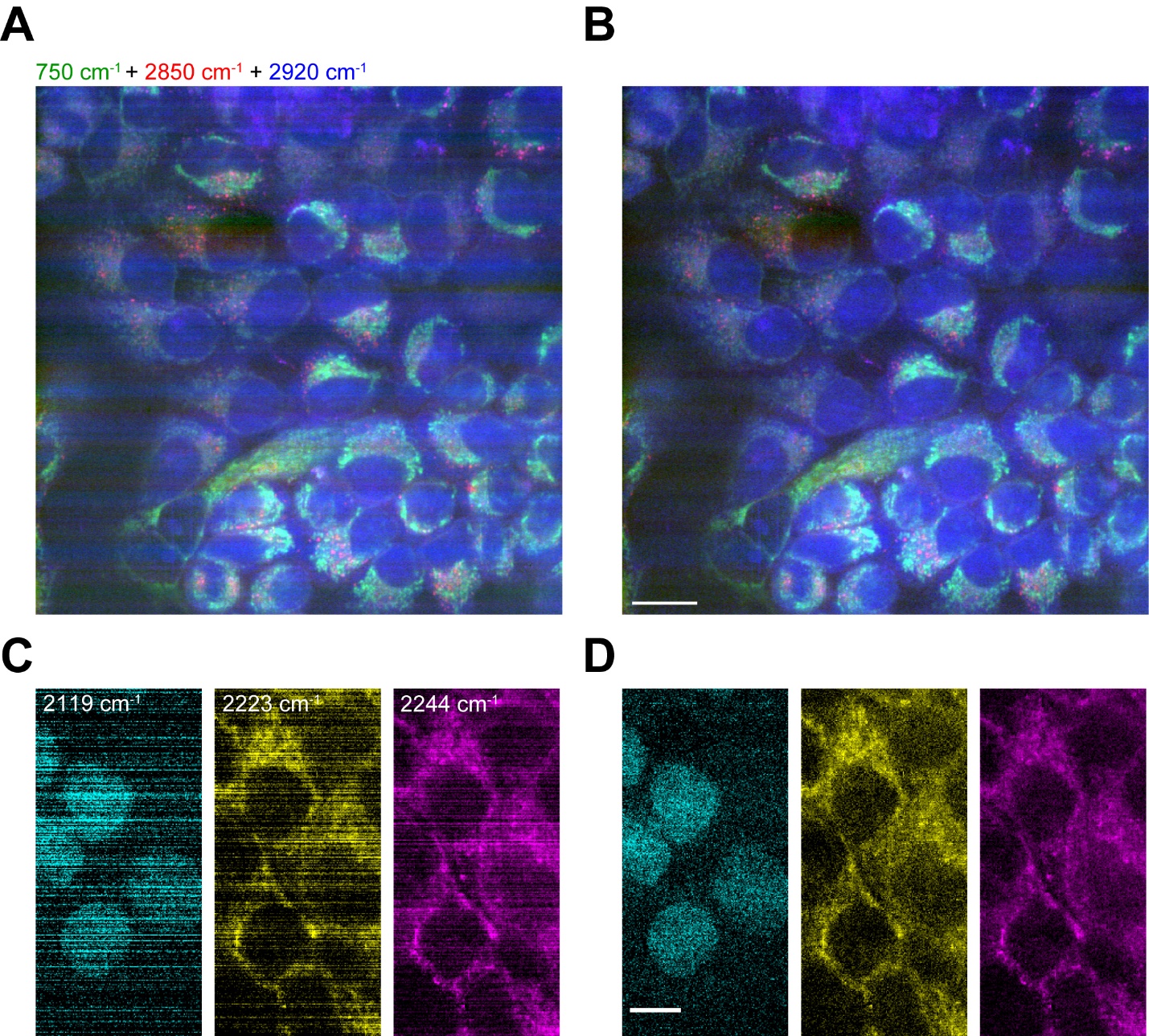

Fig. S6.

**Stripe correction of Raman images.** Raman images of cryofixed HeLa cells with wide FOV (**A**, **B**) and treated with multiple Raman tag (**C**, **D**), without stripe correction (**A**, **C**) and with stripe correction (**B**, **D**). All scale bars: 20 µm (**A**, **B**) and 10 µm (**C**, **D**)

Table S1.

**Detailed parameters for the Raman image reconstruction**

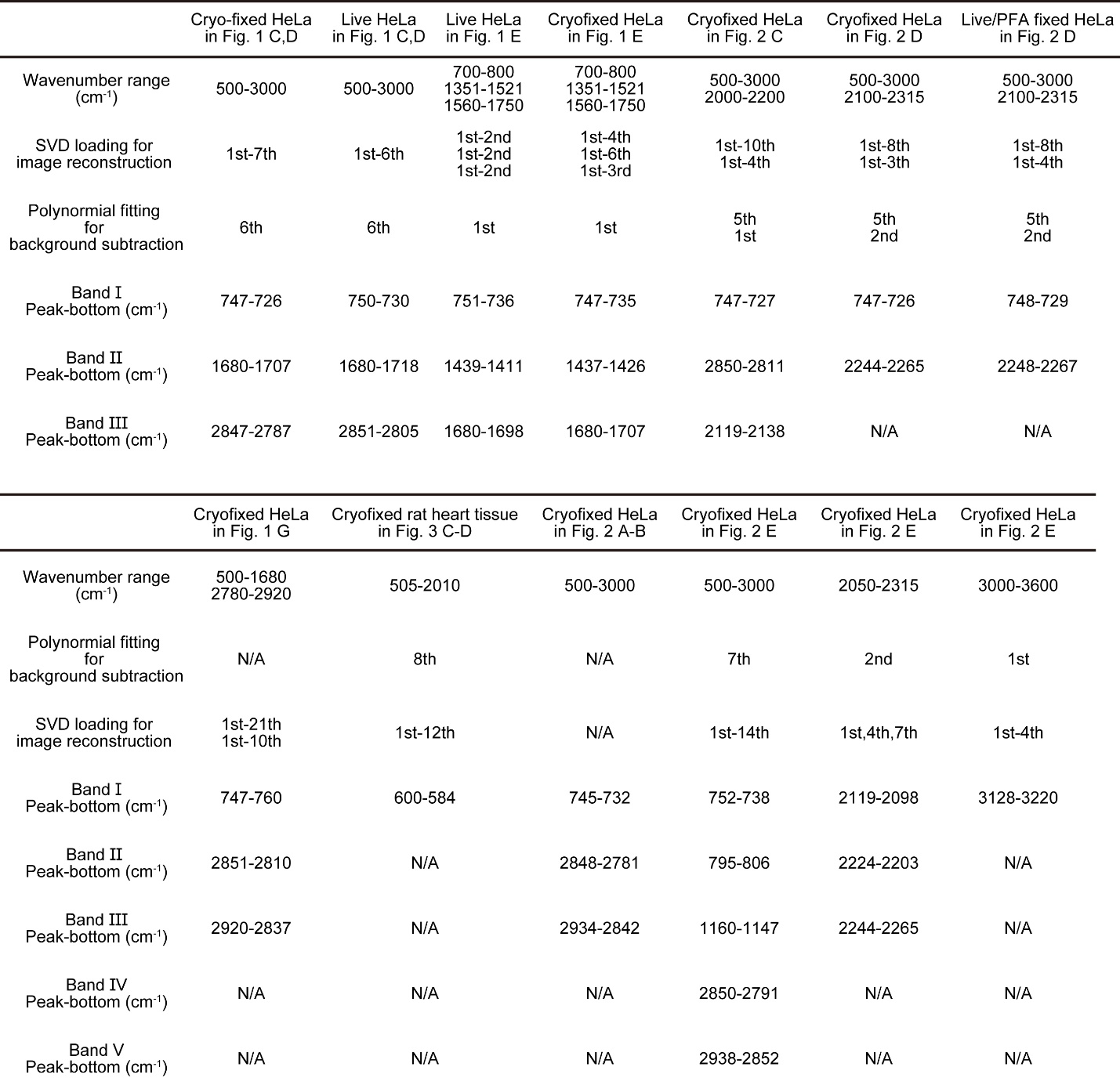
